# Streamflow drought limits fish production across river ecosystems

**DOI:** 10.64898/2026.05.05.723077

**Authors:** Timothy J. Cline, Andrew Lahr, Gregory T. Pederson, Justin Martin, David Schmetterling, Donovan A. Bell, Clint C. Muhlfeld

## Abstract

River flows sustain biodiversity, ecosystem function, and human well-being, yet climate change and water extraction are driving frequent and severe low-flow conditions (i.e., streamflow droughts). Here, we show that declining streamflow constrains fish production by limiting juvenile recruitment, the dominant demographic pathway regulating population dynamics. Across populations, declining flows nonlinearly reduce carrying capacity and production, with severe drought reducing production by up to 90%. Flow–production relationships vary among rivers, revealing both drought-sensitive and drought-resilient responses shaped by ecological and hydrological context. In some systems, maintaining higher flows during average years yields greater ecological benefits than equivalent drought-year interventions. These results demonstrate that streamflow drought directly limits biological production and highlight how adaptive water management can be implemented to sustain freshwater ecosystems under increasing hydroclimatic variability.

## Main text

River flows sustain biodiversity, ecosystem function, and critical ecosystem services for human societies (*1, 2*). However, these ecosystems face mounting threats from human use–irrigation, hydropower, municipal withdrawals–and a changing climate, including prolonged drought and rising temperatures (*3, 4*). These pressures are increasing the frequency and severity of streamflow droughts (*5, 6*): periods of extreme low flow that reshape river structure (*7*), degrade water quality (*8*), and constrain aquatic biodiversity and ecosystem function (*9-11*). Despite these growing impacts, empirical understanding of how streamflow regulates the demographic processes underpinning biological production, particularly for top consumers like fish, remains limited (*12, 13*). As streamflow droughts intensify under ongoing climate change and increasing human pressures, this knowledge gap constrains our ability to predict and manage ecosystem responses (*14*).

Freshwater fisheries–spanning commercial, subsistence, and recreational sectors–provide food, livelihoods, and well-being for millions worldwide (*15*), generating an estimated USD $122 billion annually, including $78 billion in North America alone (*16*). These benefits rely on productive and resilient fish populations that sustain biodiversity and critical ecosystem functions (*17*). However, intensifying streamflow drought poses a growing threat to freshwater ecosystems and the fisheries they underpin (*18, 19*). Prolonged low flows disrupt the timing, magnitude, and quality of streamflow regimes shaping fish life-history adaptations (*20*), while reducing habitat, elevating temperatures, lowering oxygen and food resources, and limiting access to critical spawning and rearing areas (*13*). Although numerous studies document how these stressors independently affect fish survival, growth, or reproduction (*21, 22*), the cumulative effects on fish production remain unclear. Understanding the impacts of drought on production can inform decision-making related to sustaining inland fisheries, conserving freshwater biodiversity, and securing ecosystem services that depend on healthy, functioning river ecosystems (*13*).

Here, we quantify how fish production responds to streamflow variation driven by climate variability and human water use. By linking long-term streamflow records with multi-decadal fish population data (∼1980–2023) from 25 brown (*Salmo trutta*) and 22 rainbow trout (*Oncorhynchus mykiss*) populations, we developed age-structured population models to evaluate how seasonal flows regulate juvenile recruitment, adult survival, and population production. We define fish production as the net generation of new individuals exceeding replacement, integrating demographic responses across life stages. This approach allows direct estimation of flow-dependent carrying capacity and surplus production (i.e., production of new individuals exceeding replacement), and provides a basis to evaluate how changes in streamflow translate into changes in biological production. Together, these analyses directly link hydrologic variation to population-level production, offering insights to guide adaptive water management strategies that support both ecological and societal needs.

Our study focuses on cold-water trout populations in the northern Rocky Mountains (Fig. 1a), a region with hydrologically diverse river systems that support iconic wild trout fisheries valued at over $1.4 billion annually in Montana alone (*23, 24*). Although brown trout and rainbow trout populations are non-native to the region, they have been established for over a century, are a major focus for fisheries management, and provide indicators of how streamflow constrains fish production more broadly, including for native species with similar life histories. The northern Rocky Mountains exemplify broader global water challenges, as climate warming reduces snowpack, lengthens dry seasons, and intensifies drought (*6, 25*). Concurrently, rapid development and irrigation have increased competition for water, with diversions extracting 25– 75% of midsummer streamflows in many rivers (*26*). Fisheries in this region are increasingly vulnerable as trout populations have declined in recent years, raising concern among managers, anglers, and conservationists (*27*). By linking long-term flow and fish population data across multiple species and rivers, this study provides an empirical test of how streamflow variation translates into differences in biological production in riverine ecosystems.

**Fig. 1:**
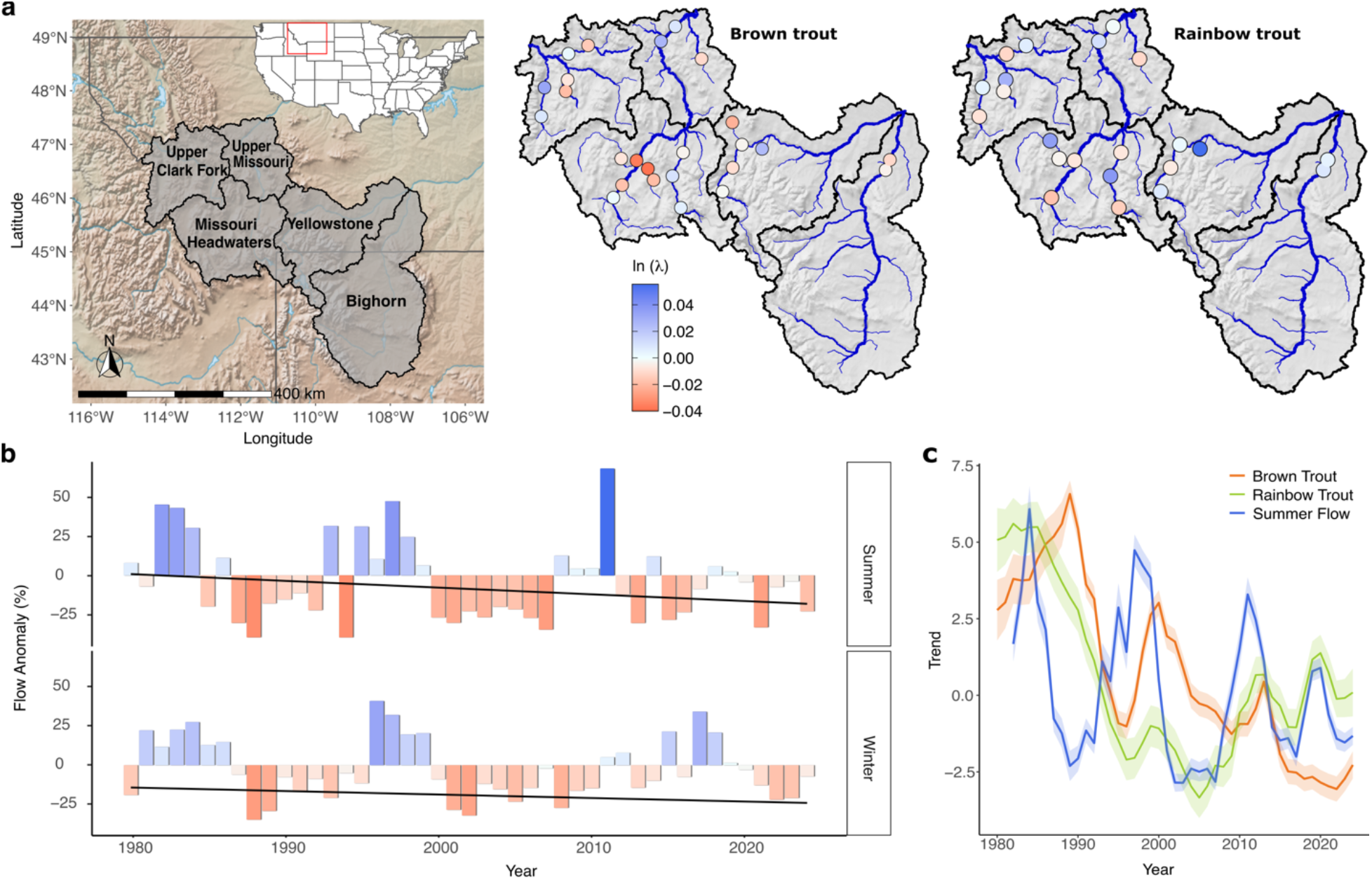
Multi-decadal declines in brown trout and rainbow trout populations closely track long-term declines in regional streamflow. (a) Map of 25 brown trout and 22 rainbow trout populations across rivers of Montana included in this study. Point colors indicate population-specific trends in abundance (red - strong decline, blue - slight increase). Population growth rates were estimated from a simple Gompertz population model (refer to Materials and Methods). (b) Aggregate summer and winter streamflow anomalies (% difference from long-term mean), based on U.S. Geological Survey (USGS) gages associated with each population, 1980-2024. Major drought periods are indicated by extended negative anomalies. Trend line was estimated using ordinary least squares regression. (c) Common trends in brown trout (brown line) and rainbow trout (green line) abundance from dynamic factor analysis of long-term mark-recapture data (1980-2024). Standardized summer streamflow trends (blue line) are overlaid for comparison. Population responses lag streamflow dynamics by 1-3 years due to life-history of each species.

### Population trends track long-term streamflow variability

Across rivers, brown and rainbow trout abundance declined markedly since 1980 (Fig. 1a-c; Table S1), coinciding with long-term regional streamflow variability (Fig. 1a-c; Table S1). These long-term declines were punctuated by episodic increases and sharp population declines that closely tracked streamflow fluctuations, with 1–3-year lags reflecting species-specific life histories (Table S2). Major declines coincided with prolonged droughts marked by multi-year negative flow anomalies (e.g., 1988-1995, 2000-2007, 2020-2025). Rebounds followed wetter periods with multi-year positive flow anomalies (e.g., 1995-1999, 2010-2011), but wet years have become increasingly rare in recent decades. Seasonal flow anomalies (Fig. 1b) revealed hydrological stress during both summer and winter, indicating the importance of evaluating summer flows for rearing and growth, as well as winter flows for overwinter survival (*28*). Together, these patterns establish a long-term linkage between streamflow variability and trout population dynamics, providing context for quantifying how seasonal streamflow influences specific demographic processes and ultimately population production in riverine ecosystems.

### Demographic responses to flow

Juvenile recruitment was consistently and positively associated with summer flows (Jul 15–Aug 31) for both species (Fig. 2a), with stronger and more consistent effects than winter flows (Dec 1–Apr 31; Fig. 2b-c). Incorporating flow covariates reduced unexplained residual variation in recruitment by 54% (Table S3). Higher summer flows expand aquatic habitat, increase food availability, and improve floodplain access, supporting juvenile survival to recruitment (*1, 29*). Flow-recruitment relationships varied among rivers, likely reflecting ecological and hydrological context, including irrigation withdrawals, dam regulation, geomorphology, thermal regime, and habitat complexity. Non-linear, quadratic responses (i.e., recruitment peaking at intermediate flows) occurred in ∼16% of populations, most commonly rainbow trout in summer and brown trout in winter, consistent with differences in spawning and rearing phenology (Fig 2a; Table S2). Rainbow trout (spring spawners) are vulnerable to egg scouring and juvenile displacement during high spring-summer flows (*28*), while brown trout (fall spawners) face similar risks during high fall-winter flows (*30*). Although summer flow emerged as the dominant driver overall, the importance of winter flow in some rivers highlights how ecological flows can be addressed across seasons based on local context. These results demonstrate the central role of streamflow in recruitment and the ecological trade-offs involved in managing flow regimes to support freshwater fish communities.

**Fig. 2:**
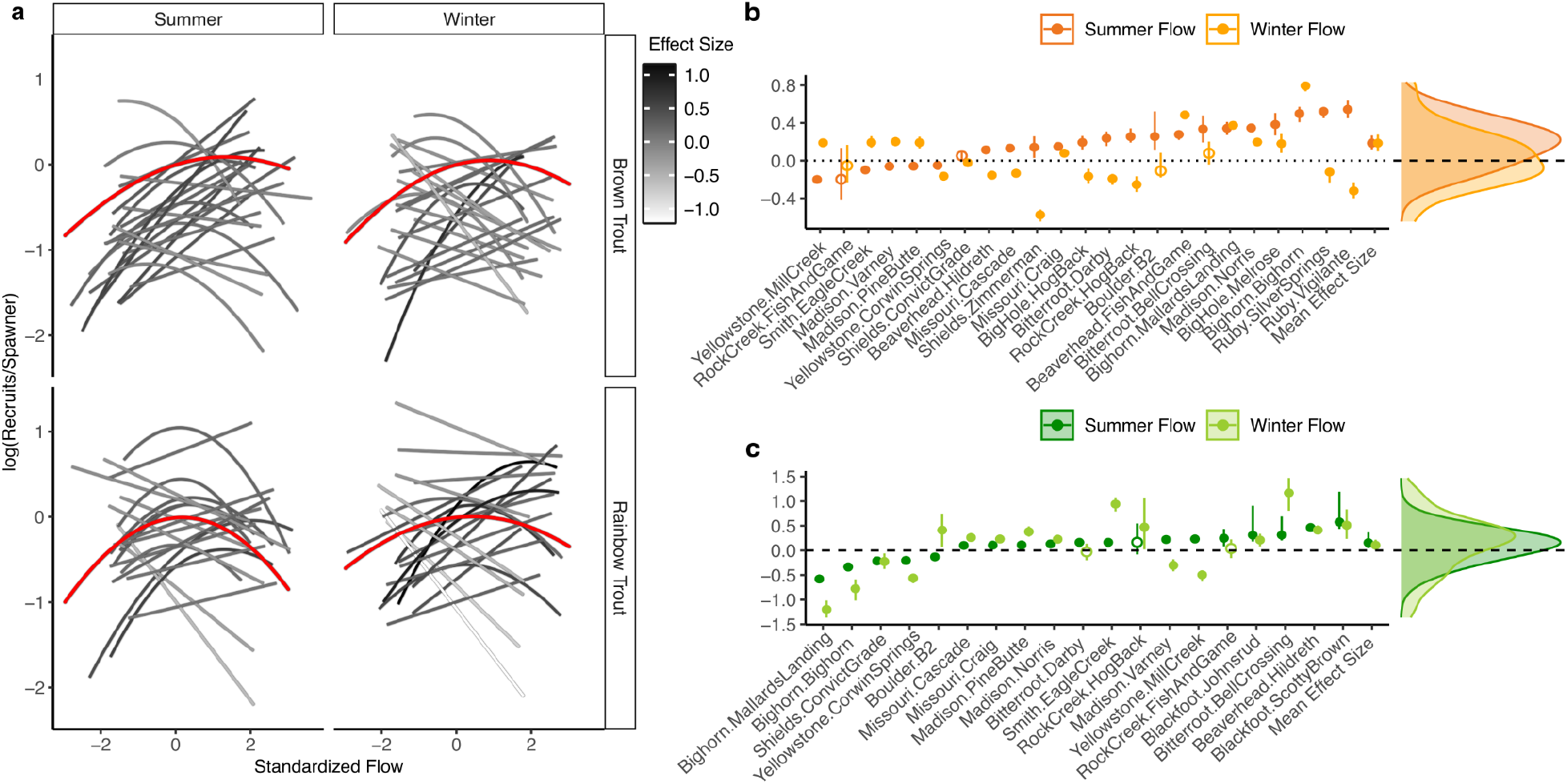
Streamflow strongly influences recruitment across brown trout and rainbow trout populations. (a) Population-specific relationships between standardized summer and winter streamflow and recruitment production [log(recruits per spawner)] for brown trout and rainbow trout populations (predicted at average spawner biomass). The red line indicates the mean effect size across populations. Streamflow covariates are lagged by 1-3 years (depending on population) to correspond to critical early life-stages (e.g., young of the year). (b,c) Median estimated effect sizes (change in y-axis value from moderate low flows, -1 SD, to median flow) and 90% credible intervals for the influence of mean summer (Jun 15-Aug 31) and mean winter (Dec 1 - Mar 31) streamflow on recruitment in (b) brown trout and (c) rainbow trout populations. Filled circles denote significant effects (credible intervals excluding zero); open circles indicate non-significant effects. Density plots (right panels) show the distribution of median effect sizes across all populations. Population labels are expressed as “river.section,” indicating river identity followed by section designation.

In contrast to recruitment, adult survival responded weakly and inconsistently to flow across populations (Fig. 3). Including flow covariates reduced unexplained residual variation in survival by only 18% (Table S3). While some populations showed modest positive or quadratic responses to flow, effect sizes were generally small, and credible intervals often overlapped zero (Fig. 3b,c). These weak effects likely reflect greater physiological tolerance and mobility of adults relative to early life stages (*13, 31*). Recruitment emerged as the dominant demographic pathway through which streamflow regulates trout production. We also found negative correlations between low summer streamflow and elevated water temperatures in systems with available temperature data (Table S4). However, streamflow consistently emerged as the primary driver of trout population dynamics in models that included both variables (Table S5). These results reinforce the importance of streamflow as the master variable regulating thermal conditions in riverine ecosystems (*1, 7, 8*). Where possible, incorporating flow-temperature interactions can improve predictions of ecological flow responses and support effective, climate-resilient strategies.

**Fig. 3:**
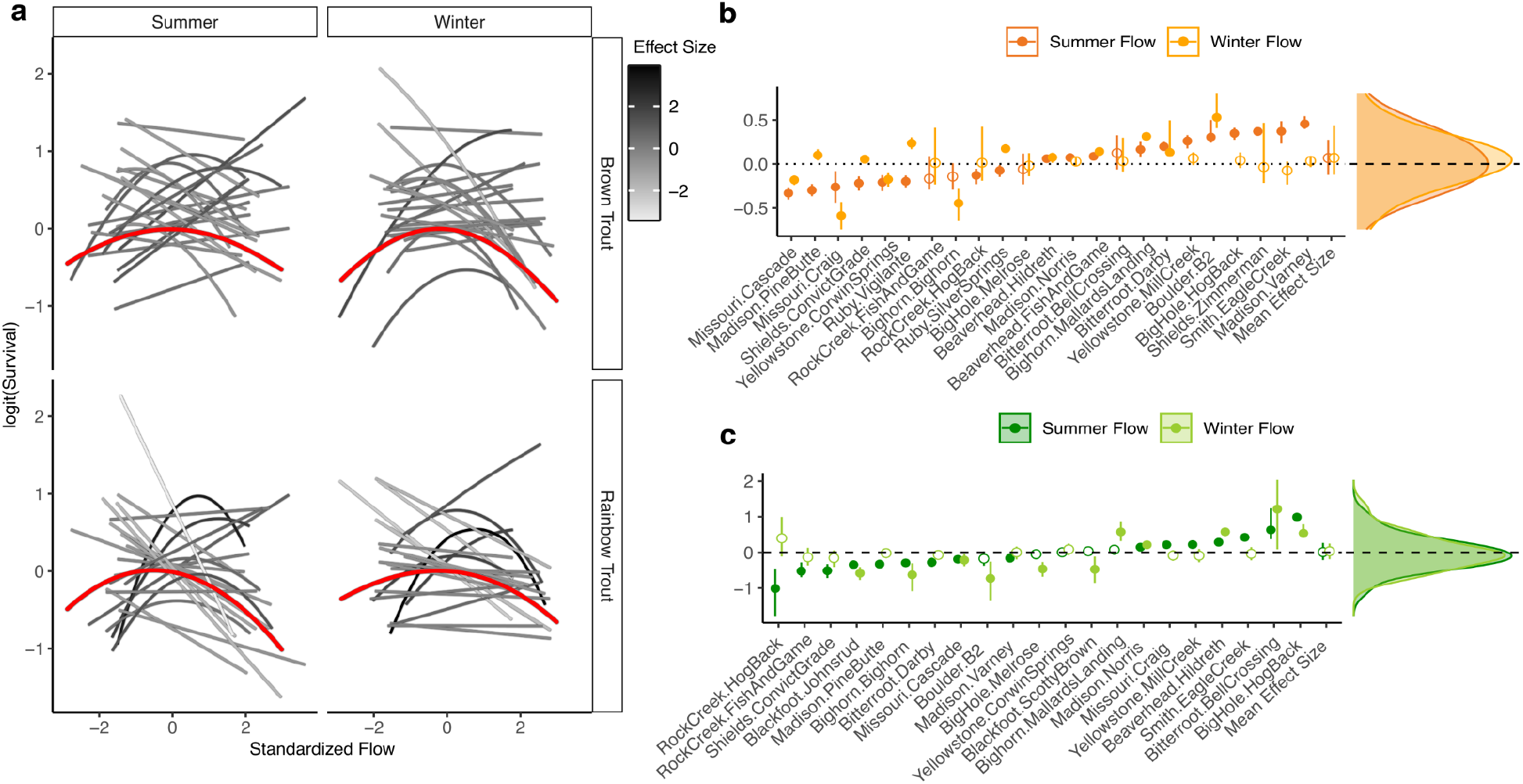
Adult survival shows weak and inconsistent relationships with streamflow across species and populations. (a) Modeled relationships between seasonal streamflow and adult survival in brown trout and rainbow trout populations. (b,c) Median estimated effect sizes (change in y-axis value from moderate low flows, where x=-1, to median flow) and 90% credible intervals for the influence of mean summer (Jul 15 - Aug 30) flow and minimum weekly average winter (Dec 1 - Apr 31) flow on survival in (b) brown trout and (c) rainbow trout populations. Filled circles denote significant effects (credible intervals excluding zero); open circles indicate non-significant effects. Density plots (right panels) show the distribution of median effect sizes across all populations. Population labels are expressed as “river.section,” indicating river identity followed by section designation.

### Flow-production dynamics

To evaluate the cumulative effects of these demographic responses on fish production, we used the fitted models to generate flow-production curves, estimating carrying capacity and surplus production across a range of hydrologic scenarios, from severe drought to wet years (Fig. 4, Fig. S1, Fig. S2). Across both species, severe drought events (5^th^ quantile, 1-in-20 year) reduced carrying capacity by 13% (IQR: -31% to 12%) and surplus production by 31% on average (IQR: -58% to 24%; Fig. 4b). Flow-production relationships varied among populations and were classified into four population response types: drought-sensitive (average surplus production reduced by >50% under severe drought), drought-resilient (reduced by 25 – 50% under severe drought), drought-resistant (little or no effect of flow), and flood-sensitive (declining production by > 25% under high flows; Fig. 4a). Drought-sensitive populations exhibited steep declines in surplus production, averaging 49% losses under moderate drought and 87% under severe drought (Fig. 4c, Fig. S2). Many of these populations are subject to substantial irrigation withdrawals (*26*), further compounding drought impacts. Drought-resilient populations also declined under drought, but retained >50% of average surplus production during moderate events, continuing to support fisheries. Drought-resistant populations showed minimal sensitivity to flow, with production largely insulated from flow variability. In regulated rivers, populations immediately below dams were often drought-resistant or resilient, buffered by cool, stable discharges that mitigated flow extremes. In contrast, populations farther downstream tended to be more sensitive to drought, where altered hydrology and water withdrawals likely intensified drought impacts. Flood-sensitive populations, predominantly rainbow trout, showed reduced production during high summer flows, likely due to species-specific differences in reproductive timing. The majority of populations, particularly for brown trout (60%), responded negatively to drought (i.e., drought-sensitive or drought-resilient; Fig. 4c). This typology provides a framework for identifying ecological flow regimes to sustain riverine fish production under increasing hydroclimatic variability.

**Fig. 4:**
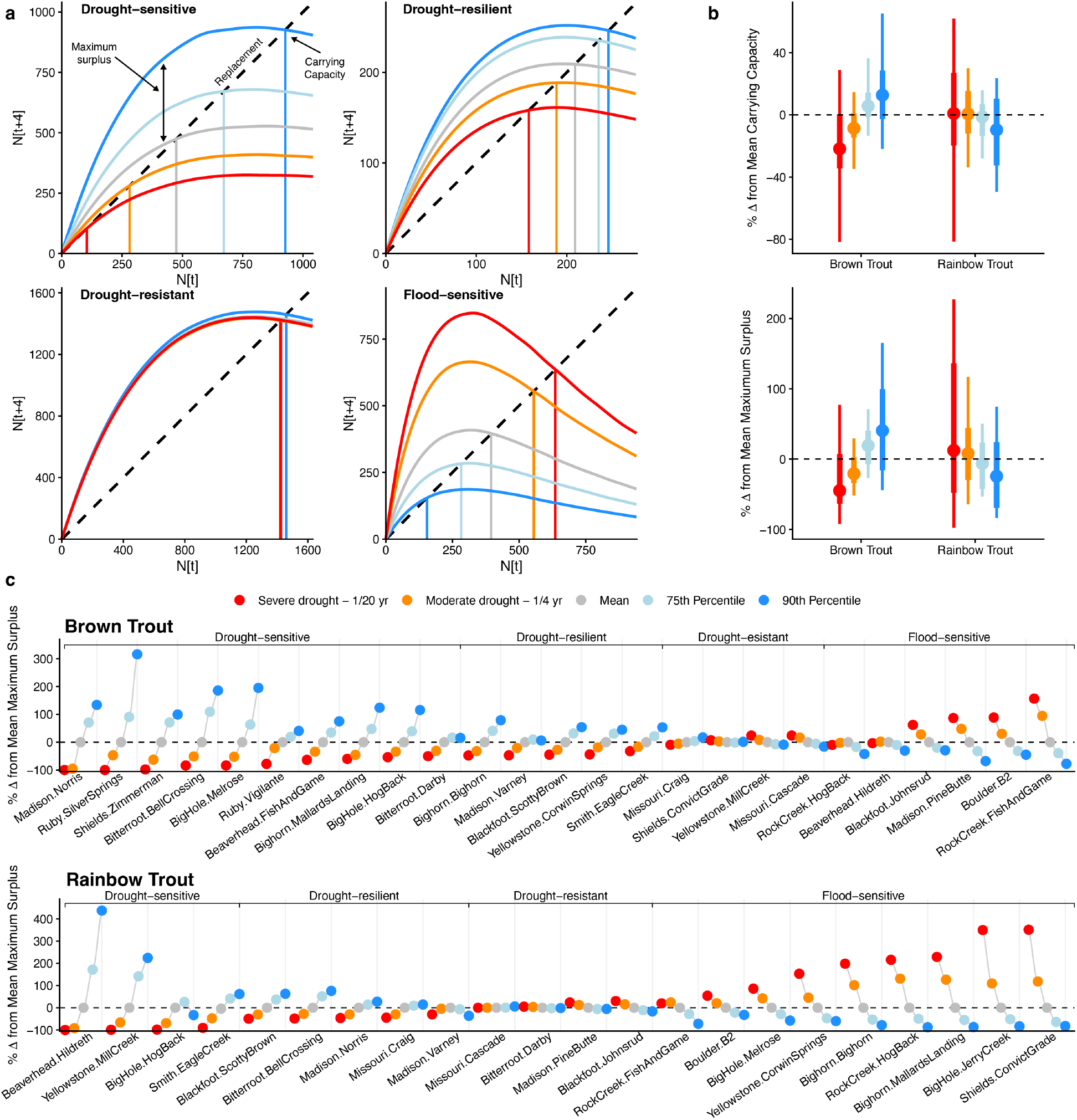
Streamflow strongly regulates trout carrying capacity and production across populations. (a) Example fish production rule curves for four representative populations, illustrating the four classifications of flow response: (i) drought-sensitive (Big Hole River, Melrose, brown trout), (ii) drought-resilient (Yellowstone River, Corwin Springs, brown trout), drought-resistant (Missouri River, Cascade, rainbow trout), and (iv) flood-sensitive (Big Horn River, Mallards Landing, rainbow trout). Each curve shows predicted production under generation interval (N_t+4_) under five summer streamflow scenarios (5th, 25th, geometric mean, 75th, and 90th percentile flows) given starting abundance N_t_. (b) Aggregate percent change in carrying capacity (top) and surplus production (bottom) relative to long-term mean scenarios across all populations under the five flow scenarios. Points indicate the median, thick bars the interquartile range, and thin lines the 10^th^ to 90^th^ quantiles. (c) Population-specific responses in surplus production (mean change from the long-term average flow scenario) for brown trout (top) and rainbow trout (bottom) populations under the five flow scenarios, shown as percent. Population labels are expressed as “river.section,” indicating river identity followed by section designation.

### Predicted flow declines and fish production

Finally, we evaluated how climate-driven reductions in summer streamflow could alter trout carrying capacity by propagating projected baseflow declines (*32*) through our population-specific flow-production models (Fig. 5; Fig. S3). Regional hydrologic projections for the upper Missouri River basin indicate that late-summer baseflows may decline by 28% by the 2080s (ensemble RCP 4.5 – RCP 8.5) (*32*). Because population-scale projections were not available for all study rivers, we implemented this decline as a standardized reduction in each population’s historical mean summer baseflow and used the resulting flow values to generate forecasts of carrying capacity and surplus production from the fitted models. Predicted responses varied strongly among populations due to heterogeneity in flow sensitivities. Under this scenario, rainbow trout capacities may decline by up 44% (10^th^ quantile) by the 2080s. Brown trout appear more sensitive, with declines up to 81% (10^th^ quantile) by the end of the century, and credible intervals indicating near-complete population loss in some systems. These reductions translate into substantial losses in carrying capacity, often exceeding the magnitude of flow declines (Fig. 5; Fig. S3). These findings indicate that, under current water management practices, climate-driven reductions in summer baseflow could translate into disproportionately large biological impacts.

**Fig. 5:**
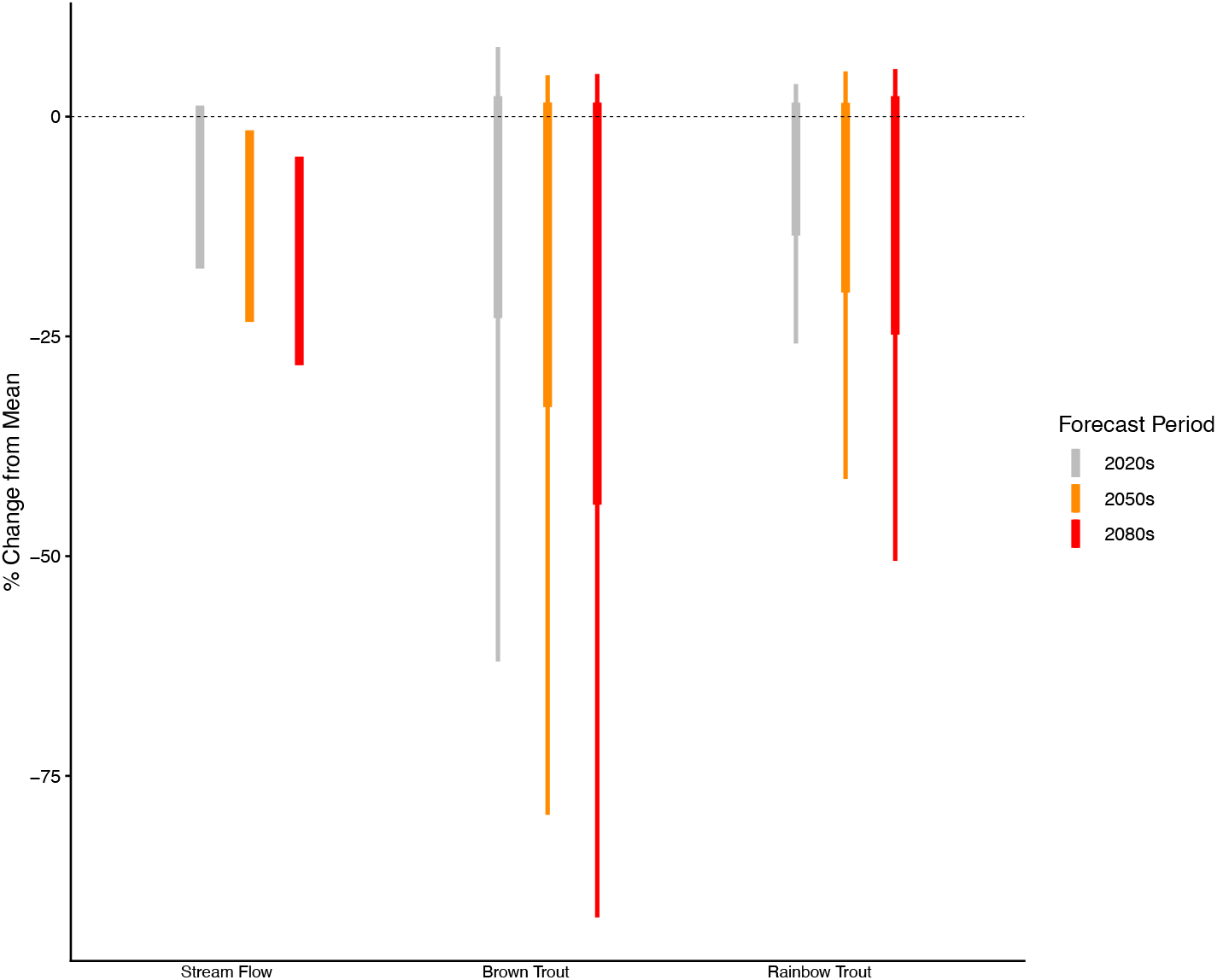
Projected future declines in streamflow are expected to substantially reduce trout carrying capacity. Percent change in carrying capacity for brown trout and rainbow trout populations under projected changes in mean summer streamflow. Projections are based on Bureau of Reclamation forecasts (*32*), showing results across a range of low and high climate-driven reductions in mean summer streamflow. Boxplots show the distribution of percent change in carrying capacity across all populations combined, with interquartile range (thick line) 10^th^ to 90^th^ quantiles (thin line).

### Ecological flows for fish production

We show that streamflow is a fundamental driver of fish production, shaping carrying capacity, resilience (i.e., surplus production), and the demographic processes that underpin riverine fishes (Fig. 1c, 2, 3, and 4b,c). Low summer flows consistently reduced recruitment and productivity, while modest increases in baseflow often yielded disproportionately large gains, particularly in drought-sensitive systems (Fig. 4a,c). These production dynamics likely reflect drought-driven constraints on both adult and juvenile life stages: reduced growth and reproductive output in spawning adults (*33*), intensified competition for limited spawning habitat (*34*), and diminished juvenile growth during periods of high energetic demand, which lowers overwinter survival (*35*) (Table S2). While extensive research has documented how drought alters freshwater habitats (*7*), metabolism (*36*), and community structure (*10*), our findings provide rare empirical evidence that streamflow drought limits biological production at the ecosystem scale, integrating effects across life stages, populations, and time. Recognizing these dynamics highlights how ecological flow assessments can advance beyond habitat surrogates (*37*) and correlative models (*38*), toward mechanistic, empirically grounded frameworks that link streamflow to demographic and ecological processes (*14*). The framework developed here offers a transferable basis for linking hydrologic changes to biological production, with broad relevance for managing fisheries and understanding ecosystem responses across riverine food webs.

Forward-looking management approaches that integrate both ecological processes and human needs (*39, 40*) can support conservation of freshwater ecosystems under growing water scarcity. The ongoing 2020–2025 drought in the northern Rocky Mountains illustrates how climate-driven reductions, compounded by irrigation withdrawals, are already constraining fish population recovery and biological production in many rivers (Fig. 1b). Current minimum flow thresholds– often set at the 5th-10th percentile–may be insufficient to sustain fish productivity under increasingly extreme and variable hydrologic regimes (*41, 42*). Maintaining higher flows during average and wet years may deliver greater ecological benefits than allocating scarce water during drought, when biological responses are diminished (Fig. 4a,c) and tradeoffs with human use are most acute. Such a shift could help further incorporate resilience-building and climate adaptation into water and fisheries management. Maintaining higher flows also reduces stream temperatures and thermal stress (*43*), which could further reduce drought impacts on fish. A range of innovative and flexible water management tools–such as instream flow leasing, rotational fallowing, water banking, aquifer recharge, environmental water markets, and adaptive hydropower operations–offer scalable solutions to support ecological flows while balancing competing demands (*44*).

### The value of water for resilient fisheries

Drought resilience in river ecosystems depends not only on biological productivity but also on the capacity of human communities to adapt to changing water conditions, analogous to resilience mechanisms in other coupled social-ecological systems (*45*). During periods of low flow and elevated temperatures, fishers shift their efforts from drought-sensitive to drought-resilient rivers, stabilizing regional participation and economic returns (*23*). Because flow regimes influence both fish production and the regional economies they support, drought-resilient rivers that sustain trout productivity under low-flow conditions serve as critical ecological refuges and socioeconomic buffers. Anticipating population responses to streamflow drought requires identifying vulnerabilities within and among rivers, which can inform conservation of valuable or at-risk species and support more targeted and coordinated management, such as setting instream flow targets or adaptive fishing regulations, across heterogeneous landscapes.

While water’s value is well-established in agriculture, energy, and municipal sectors, its importance for sustaining economically and ecologically vital fisheries has been largely overlooked (*46*). Although full restoration of natural flow regimes is rarely feasible (*1, 39*), ecologically informed strategies can help design alternatives that balance environmental and human needs (*40, 41*). By quantifying how streamflow translates to fish population gains, this work provides a foundation for how ecological production could be incorporated into water allocation decisions. Ecological flows can be integrated into water policy and management to sustain biodiversity, ecosystem services, and the resilience of freshwater fisheries under accelerating environmental change (*47*).

## Acknowledgements

This work was supported by the National Oceanic and Atmospheric Administration (NOAA) Climate Program Office National Integrated Drought Information System and the U.S. Geological Survey (USGS) Ecosystems Mission Area’s Ecological Drought Program. We thank Britt Parker (NOAA-NIDIS) for project management and the many biologists from Montana Fish, Wildlife & Parks who collected the long-term trout population data. We also thank Joe Giersch for assistance with final graphics. Any use of trade, firm, or product names is for descriptive purposes only and does not imply endorsement by the U.S. Government.

## Author Contributions

Conceptualization: TJC, CCM, GP, JM, DS, AL, JM, DB

Methodology: AL, TJC, CCM, DB

Investigation: TJC, CCM, GP, JM, DS, AL, JM, DB

Visualization: AL, TJC, CCM, DB

Funding Acquisition: TJC, CCM, GP

Writing – original draft: TJC

Writing – review & editing: TJC, CCM, GP, JM, DS, AL, JM, DB

## Data, code, and materials availability

All data and code to support this study is available on GitHub at https://github.com/andrewlahr/FlowFishProduction.

## Supplementary Materials

Materials and Methods

Figs. S1 to S3

Tables S1 to S5

## Material and Methods

### Data

We used trout population monitoring data from 1980 to 2023 across 25 reaches along 13 rivers (5th to 7th order) in Montana, totaling 720 surveys along 128 river kilometers ^1^. Data were collected by Montana Fish, Wildlife & Parks as part of annual-to-semi-annual, closed population capture-mark-recapture electrofishing surveys, which included fish species, weight, length, marking identification, and section length. During the marking run, all fish received a fin clip for identification during the subsequent recapture run. To ensure ample variation within the data and improve model performance, time series needed to span at least 20 years and have a maximum of 5 years between surveys. We limited data to post-1980 to remove the effects of stocking, which ceased in Montana rivers in the late 1970s.

Daily streamflow data were acquired from U.S. Geological Survey (USGS) streamgages located near each fisheries survey section using gages utitlized by the state for making reach specific management decisions^2^. Continuous discharge readings were summarized into summer (Jul. 15-Sep. 30), and winter (Dec. 1-Mar. 30) means. Flow metrics were log transformed, centered, and scaled. Temperature data were only available at approximately half of the sites and often had substantially shorter record lengths than streamflow, limiting their utility. However, where available, they were often correlated with mean summer streamflow (Table S4).

## Analysis

### Abundance estimation

Age-specific observed abundance time series were estimated using closed-population mark recapture data, survey section length, age estimates, and log-likelihood abundance estimators. To be consistent with Montana Fish Wildlife and Parks, our abundance estimation procedures used the following approach ^3^. Abundance was estimated using a mark-recapture methodology that corrected for variation in detection probability based on fish length. Fish were first grouped into 25 mm length bins (*length*.*bin*). The midpoint of the bin was used as the length covariate in the regression analyses (e.g., minimum bin length plus 12.5 mm).

Next, we modeled whether fish *i* caught during the recapture run was clipped (i.e., also caught during the marking run). This was treated as a binary outcome with *y*_*i*_ *=* 0 for fish that were not clipped and *y*_*i*_ *=* 1 for fish that were clipped. This binary outcome was modeled using a generalized linear model (GLM) with a Bernoulli distribution and a logit link:d

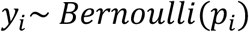

where *p*_*i*_ represents the detection probability for fish *i. p*_*i*_ was modeled using four equations that test different hypotheses about the relationship between fish length and detection probability:

1. Null model: *logit*(*p*_*i*_) = *β*_0_
2. Length linear: *logit*(*p*_*i*_) = *β*_0_ + *β*_1_*length.bin*_*i*_
3. Length quadratic 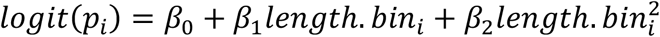
4. Length quadratic fixed intercept: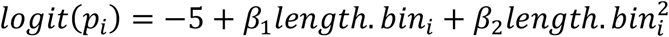

We calculated AICs ^4^ for each model. We then predicted detection probability (*p*_*l*_) for each length bin *l* using weighted model averages.

To estimate total abundance, we next summed the abundance estimates for each length bin. Specifically, the number of fish caught during the mark run in each length bin was tabulated (*C*_*l*_) and divided this by the length bin specific detection probability to get a length bin specific estimate of abundance. These estimates were summed from bin 1 to the maximum bin *L*:

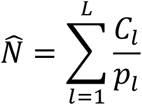

The standard error of 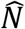 was calculated by first using the delta method to propagate uncertainty from *p*_*l*_ to the length bin specific abundance estimate. We then calculated the standard error of total abundance estimate by taking the square root of the sum of the squared standard errors of the length-specific abundance estimates. Population estimates were then divided by the length of the survey section to standardize abundance to distance sampled (fish/km). Log-likelihood abundance estimators were run in the R Statistical Programming Environment ^5^.

### Population trend analyses

To estimate population growth rate (lambda) for all populations in our analysis (displayed only in Figure 1), we estimated a population growth rate parameter (*r* or log(*lambda*)) for each population individually (Fig. 1a), using a simple Gompertz population model,

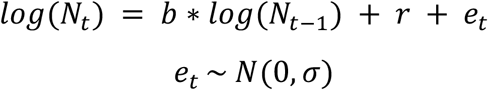

Where *N*_t_ is the population abundance in year *t, b* represents negative density-dependence, and *r* is the population growth rate over the time series. We fit these models using the MARSS package ^6^ in R. Population growth rate estimates for each population are presented in Fig. 1a and available in Table S1.

To evaluate general trends in trout abundance across rivers in Montana, we used Dynamic Factor Analysis (DFA,^7^, a dimension reduction technique that summarizes common trends among many time series (Fig. 1c). With DFA, we are trying to explain temporal variation in a set of *n* observed time series using linear combinations of independent hidden random walks. The model structure is as follows:

The state equation of a vector common random walk trends over time:

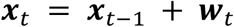

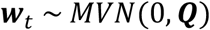

Observation equation relates trend (***x***) to observations (***y***).

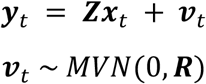

Here, the vector of observations for abundances in each population at time *t* (***y***_t_) are modeled as linear combinations of hidden trend (***x***_t_) and factor loadings on the hidden trend for each population (***Z***). 𝒱_t_ and *w*_t_ represent the observation and process error structures, respectively. To make the model estimable^6^ process error (***Q***) was set to a diagonal matrix of value 1. Observation errors (***R***) are from a multivariate normal distribution. Since we were only interested in the single most parsimonious trend among populations, we only fit DFA models allowing 1 common trend. We compared models with two observation error structures; where all stocks have individual observation error (diagonal and equal) and where populations have individual observation error (diagonal and unequal). Candidate models were compared using AIC based on the maximum likelihood of the model fit^3^. We fit an independent single-trend DFA analysis to 1) all rainbow trout populations and, 2) all brown trout populations (Fig. 1c). All data were z-scored prior to analysis to account for differences in means and intrinsic variance dynamics. DFA models were fit using the ‘MARSS’ package in R ^6^. Trend loading parameter estimates for each population are available in Table S1.

### Aging and abundance estimates

We used Bayesian hierarchical Gaussian mixture models to estimate the mean size-at-age and residual error for three age-classes (Age 2 or recruits, Age 3, Age 4+) in each population using annual length frequency data.

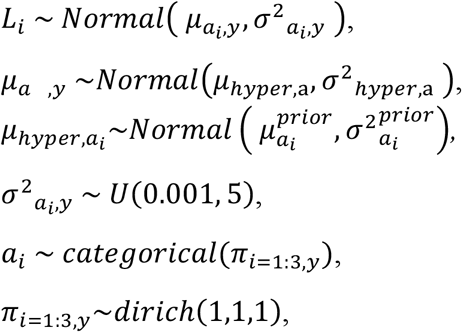

Where *L*_*i*_ is the length of individual *i*,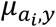 is mean size at age of a given age class *a*_*i*_ in year *y, σ*^2^_*a,y*_ is the residual variance for mean size at age of age *a* in year *y*,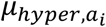 is the hyper prior used to inform mean size of age class *a*_*i*_ with residual variance 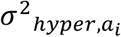. We leveraged existing size-at-age data from previously conducted otolith and scale analyses by Montana Fish, Wildlife, & Parks at a small sample of populations and expert opinion from local biologists to determine 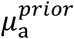 or site-specific and statewide mean size at age, and maximum and minimum bounds on µ_*a,y*_ (*L*_*min,a*_, *L*_*max,a*_). Hierarchical Gaussian-Mixture models were completed in the JAGS statistical program ^8^, ran in the R Statistical Programming Environment ^5^ using the rjags ^9^ and R2Jags ^10^ package. We used a burn-in of 1500 iterations, ran 6000 additional iterations, thinned the chains by 10, and included 3 chains. All models were run to convergence as determined by the potential scale reduction factor, R-hat <1.1 for all parameters ^11^

### Population Models

We developed age-structure integrated population models to investigate the influence of streamflow on the dynamics of trout populations. The population abundance (fish/km) for each age class each year is described as,

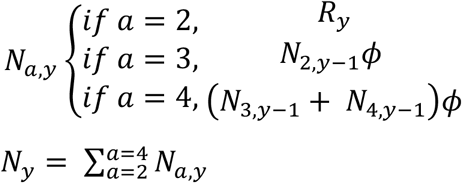

where *N*_*a,y*_ is the abundance of age-class, *a*, during year *y, R*_*y*_ is the abundance of recruits in year *y*, and ϕ_*y*_ is the survival of fish beyond recruitment in year *y*. We assume recruitment occurs at age 2 and spawning begins at age 3 based on otolith analysis and consultation with regional biologists. Maximum age was limited to 4+, as mixture models could not sufficiently differentiate beyond three age classes, and all fish over age 2 are assumed to be reproductive.

Recruitment was estimated using a log-linear Ricker stock-recruitment relationship ^12^, influenced by spawning stock biomass and mean seasonal flow,

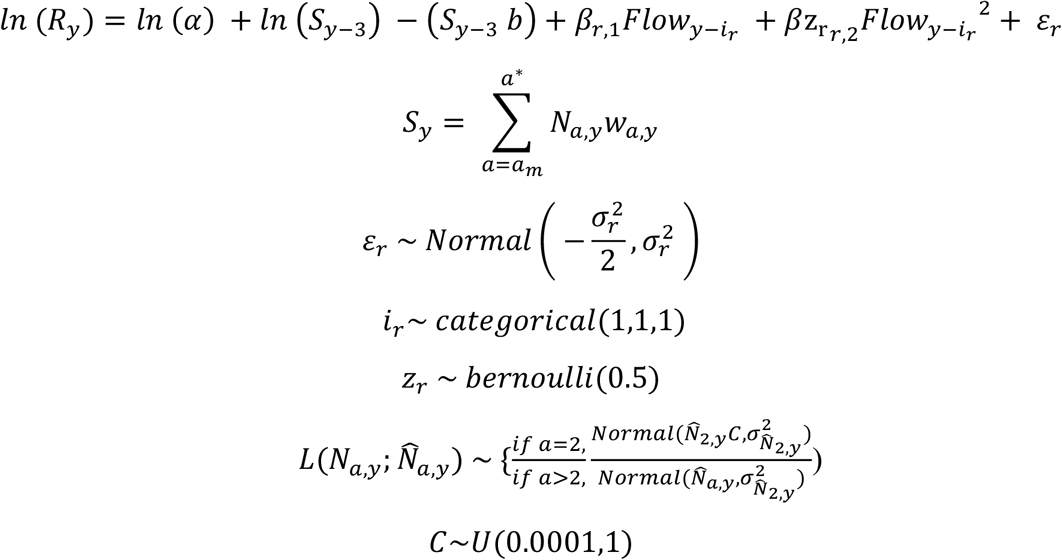

where ***S***_*y*-3_ is the biomass abundance (units) of the stock that produced year *y* recruits, *i*_*r*_ is the lag between flow experienced during the unobserved years between spawning and recruitment, 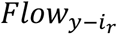 is the standardized mean seasonal flow experienced during recruitment, *a*_*m*_ is age at sexual maturity or minimum age of spawners, *a*^∗^ is maximum age, *w*_*a,y*_ is the mean weight of age class *a* in year *y* from survey data, and 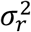 is the variance of log recruitment. Parameters *α* and *b* are estimated Ricker model parameters and *β*_*r*,1_ and *β*_*r*,2_ are the estimated linear and quadratic effects of mean seasonal flow on recruitment. To account for lower catchability of small (<150 mm) fish in large river electrofishing, we incorporated a catchability parameter, C, within the likelihood statement for recruit abundance. We tested models with varying time lags between stream flow and recruitment, *i*_*r*_, to investigate how flow influences different stages of recruitment. Lags varied depending on the season of the flow in the model and stage of interest (Table 1). A categorical prior was set on *i*_*r*_ and a Bernoulli-distributed dummy variable *z*_*r*_ was incorporated to assess the inclusion of each lag and quadratic terms.

**Table 1.**
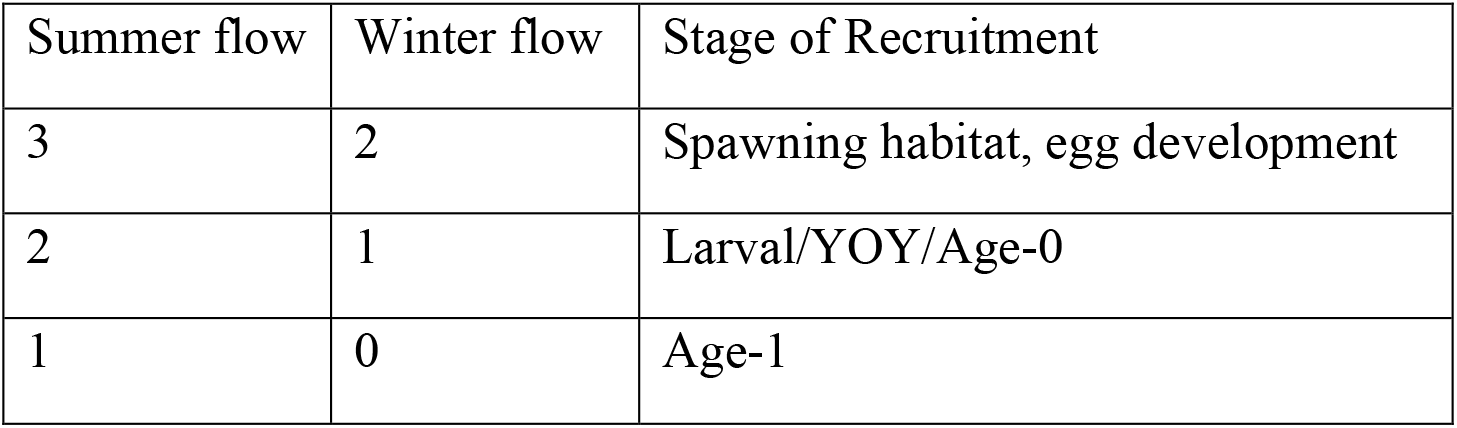
Description of lags between streamflow and different stages of recruitment.

Annual survival probability was modeled as log-normally distributed around a mean survival (ϕ_0_) and population abundance in the previous year (*N*_*y*-1_) and lagged streamflow, with a standard deviation 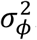. A Bernoulli-distributed dummy variable (*z*_*s*_) was incorporated to assess the inclusion of each lag and quadratic terms.

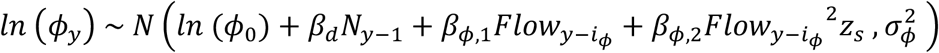

Like recruitment, the lag between flow and survival, *i*_ϕ_, depended on the season of flow modeled and the timing of population sampling. For populations sampled in the spring, mean summer flow was lagged by one year, and mean winter flow was lagged by zero years; for populations sampled in the fall, both summer and winter flow metrics were lagged by zero years. When mean summer and winter flow were correlated (p > 0.5, Table S1) we ran separate univariate analyses for recruitment and survival to avoid overfitting; otherwise, global models were used.

All integrated population models were completed in a Bayesian framework using the JAGS statistical program, run in the R statistical programming environment using the rjags and R2Jags packages ^5,8-10^. We used a burn-in of 20,000 iterations, ran 80,000 additional iterations, thinned the chains by 10, and included 4 chains. All models were run to convergence as determined by the potential scale reduction factor (R-hat) being less than 1.1 for all parameters following ^11^. Uninformative uniform priors were used for Ricker model parameters, as there is limited information on stock-recruitment relationships for inland trout. The density-dependent Ricker model parameter *b* is typically small and < 0, so priors were truncated between 0 and 0.0001, while values for *α* are not often reported. Normally distributed priors on a logit scale, centered around 0 with a standard deviation of 3, were set for *ϕ*_0_ and all linear recruitment and survival covariate effects, while priors for quadratic covariate effects were constrained to be less than zero, as the relationship between streamflow and fish production is typically concave.

## Model selection

We used stability selection ^13^ to fit the most appropriate lag and covariate structure for each population. Within this framework, models cycled through recruitment-flow lags while simultaneously turning on and off quadratic terms with each iteration, ultimately converging on the most optimized set of lags and terms for a given population. To quantify the inclusion of each lag, a categorical prior was set on *i*_1_ that allowed the model to change the time lag with each iteration. Simultaneously, the Bernoulli-distributed dummy variables, *z*_*r*_ and *z*_*s*_, were used to assess the inclusion of the quadratic terms, *β*_*r*,2_ and *β*_*ϕ*,2_. The number of lag and quadratic model combinations varied depending on whether a global or univariate model was used. Global models had nine possible summer and winter flow lag pairs and sixteen different quadratic inclusion outcomes for each, totaling 144 possible models, while univariate linear models had three lag combinations with four different quadratic outcomes for each season, totalling 24 models. To ensure ample time for convergence, we used a burn-in of 187,500 iterations; models ran for an additional 750,000 iterations, thinned the chains by 10, and included 4 chains.

Separate models, described for population, were run with selected (inclusion probability > 0.5) lag and covariate structure to optimize parameter estimation (Table S2).

We evaluated the strength of flow effects on recruitment and survival using posterior distributions of covariate coefficients and associated credible intervals. For each demographic process, we quantified explanatory power as the proportional reduction in residual process variance,

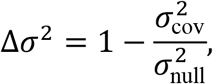

computed independently for recruitment and survival time series by comparing models with flow effects to otherwise identical models excluding covariates. These variance reduction values are available in Table S3.

### Stream Temperature

Where temperature data was available and was not highly correlated streamflow, we employed a similar stability selection framework to assess the inclusion of summer stream temperature in our analysis. To quantify the inclusion of temperature variables, we parameterized models with additional quadratic temperature effects on recruitment; *β*_*rt*_, *β*_*rt*2_ and survival; *β*_*ϕt*_,*β*_*ϕt*,2_ and paired Bernoulli distributed dummy variables; *z*_*rt*_ and *z*_*st*_. If the median value of either dummy variable’s inclusion probability was greater than 0.5 in each population, temperature was an influential factor in demographic processes in addition to stream flow. Of the 13 sites with stream temperature data, 2 showed strong (Pearson’s R >0.5) collinearity between summer temperatures and streamflow and were therefore excluded from this analysis, leaving 11 rainbow trout and 11 brown trout populations. Of these, no brown trout populations and only two Rainbow trout populations had inclusion probabilities greater than 0.5 on either dummy variable (Table S4). It should be noted that temperature data time series were often shorter than those of streamflow, limiting direct inference in final model structures.

### Simulation models

Streamflow-fish production rule curves were generated by simulating each population under constant flow for one generation (4 years), using 1,000 randomly drawn parameter sets from posterior parameter distribution of the integrated population models. Each parameter set was used to simulate populations with a range of initial stock abundances (*N*_*t*_) from 1 to 5,000 under five mean summer flow quantiles: 5^th^, 25^th^, 75^th^, and 90^th^ percentiles, as well as the geometric mean flow. The resulting population sizes were projected to the next generation (*N*_*t+4*_). We constructed each rule curve and its variance by calculating the interquartile range of each *N*_*t+4*_ value across parameter sets for each flow scenario. Each set of *N*_*t+4*_ and *N*_*t*_ values was smoothed using spline functions to reduce noise for visualization purposes. We identified the carrying capacity as the value of *N*_*t+4*_ at which *N*_*t+4*_ *-N*_*t*_ was minimized, and maximum surplus production as the maximum value of the difference between *N*_*t*_ and *N*_*t+4*_ for each parameter set and flow scenario. We then calculated the interquartile range of carrying capacity and maximum surplus production across parameter sets to obtain each value for each flow scenario. Streamflow-fish production rule curves were simulated using the R statistical programming environment.

### Future streamflow projections

To estimate the impact of projected future streamflow conditions on future fish populations we utilized estimates of future streamflow developed by the Bureau of Reclamation for the Missouri Headwaters Basin Study ^14^ and detailed in the Upper Missouri Basin Impacts Assessment: Water Supply ^15^. These future streamflow estimates were produced using downscaled Global Climate Model (GCM) – based projections of future climate conditions fed through the Variable Infiltration Capacity (VIC) hydrology model ^16,17^. GCM projections used in this effort included a subset of both CMIP3 ^18^ and CMIP5 ^19^ projections representing greenhouse gas (GHG) emissions scenarios CMIP3 A1B, CMIP5 RCP 4.5, and CMIP5 RCP 8.5 which resulted in an ensemble of 216 individual projections representing a range of plausible future GHG emissions scenarios. All GCM projections were downscaled using the Bias Correction and Spatial Downscaling (BCSD) approach ^20^ and future climate projections for use in VIC were developed using the ensemble-informed hybrid delta (HDe) method ^21,22^.The HDe method results in a time series of projected temperature and precipitation that captures the change in future climate projected by the GCM’s while maintaining spatial and temporal patterning present in the observational climate data for the region. Finally, the ensemble of projections was subset into three future time periods made up of 30-year daily time-series centered on the years, 2020, 2050, and 2080. These data then served as model inputs to the VIC model on a 1/8^th^ degree grid. Streamflow percent changes from the historical period (1950-1999) were averaged across the Headwaters region of the Upper Missouri River Basin (subbasins upstream of Canyon Ferry Reservoir) for each of the three future time periods and the range of future hydrologic projections resulting from the full ensemble of future climate scenarios was used to define the maximum and minimum plausible change in future streamflow for each period. Because these projections are not stream specific, we applied a constant percent reduction in baseflow based on the above projections to each populations historical mean flow. To create rule curves for summer flows under future drought forecasts, we applied the lower and upper bounds of forecasted percentile change in mean flow under each forecast period (2020s, 2050s, 2080s), providing six additional flow scenarios specific to each river.

