## Supplemental Figures for "Streamflow drought limits fish production across river ecosystems"

Supplementary Figures

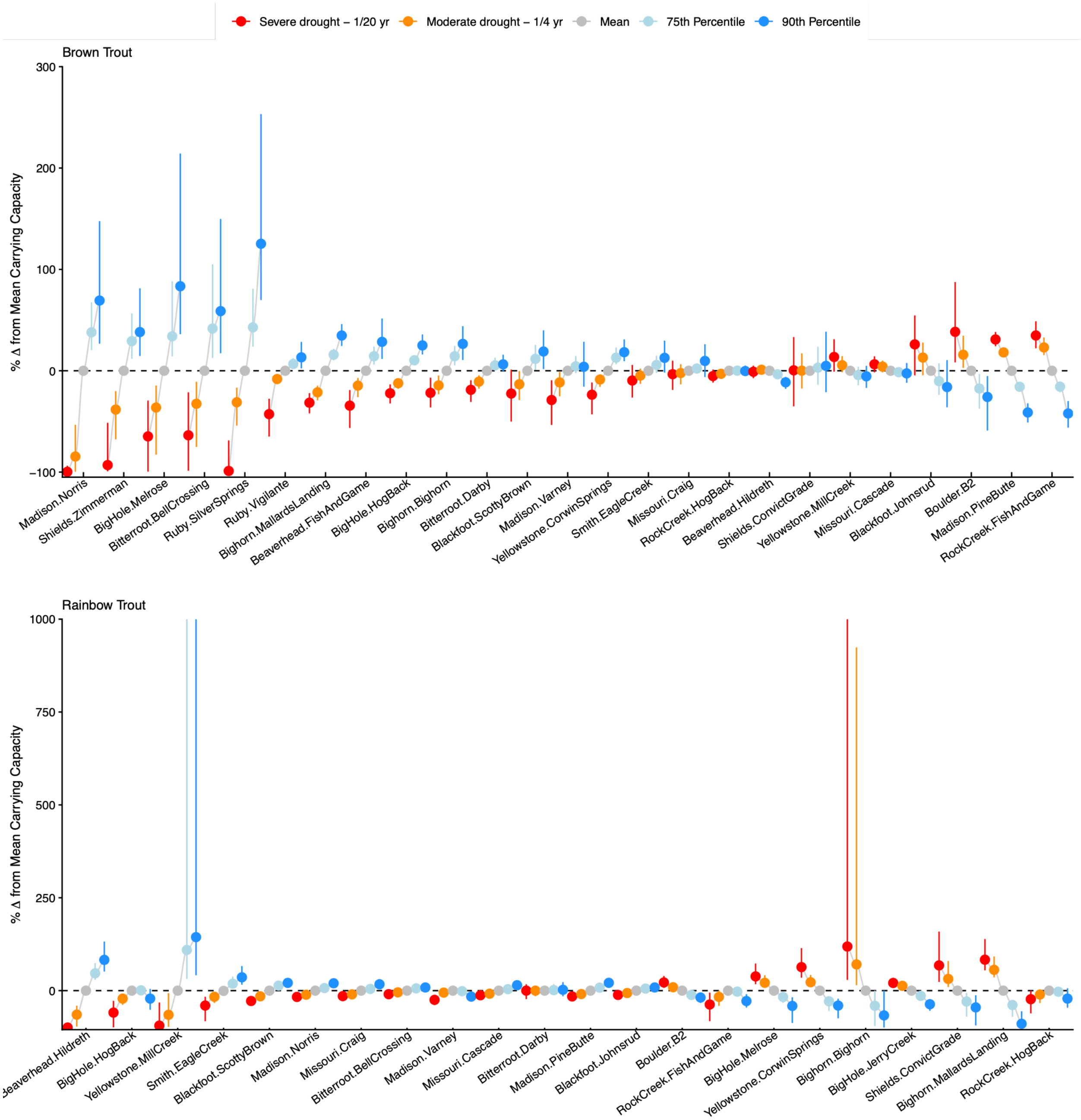


Fig S1. Population-specific responses in carrying capacity (mean change from the long-term average flow scenario with 90% credible intervals) for brown trout (top) and rainbow trout (bottom) populations under the five flow scenarios, shown as percent. Population labels are expressed as “river.section,” indicating river identity followed by section designation.

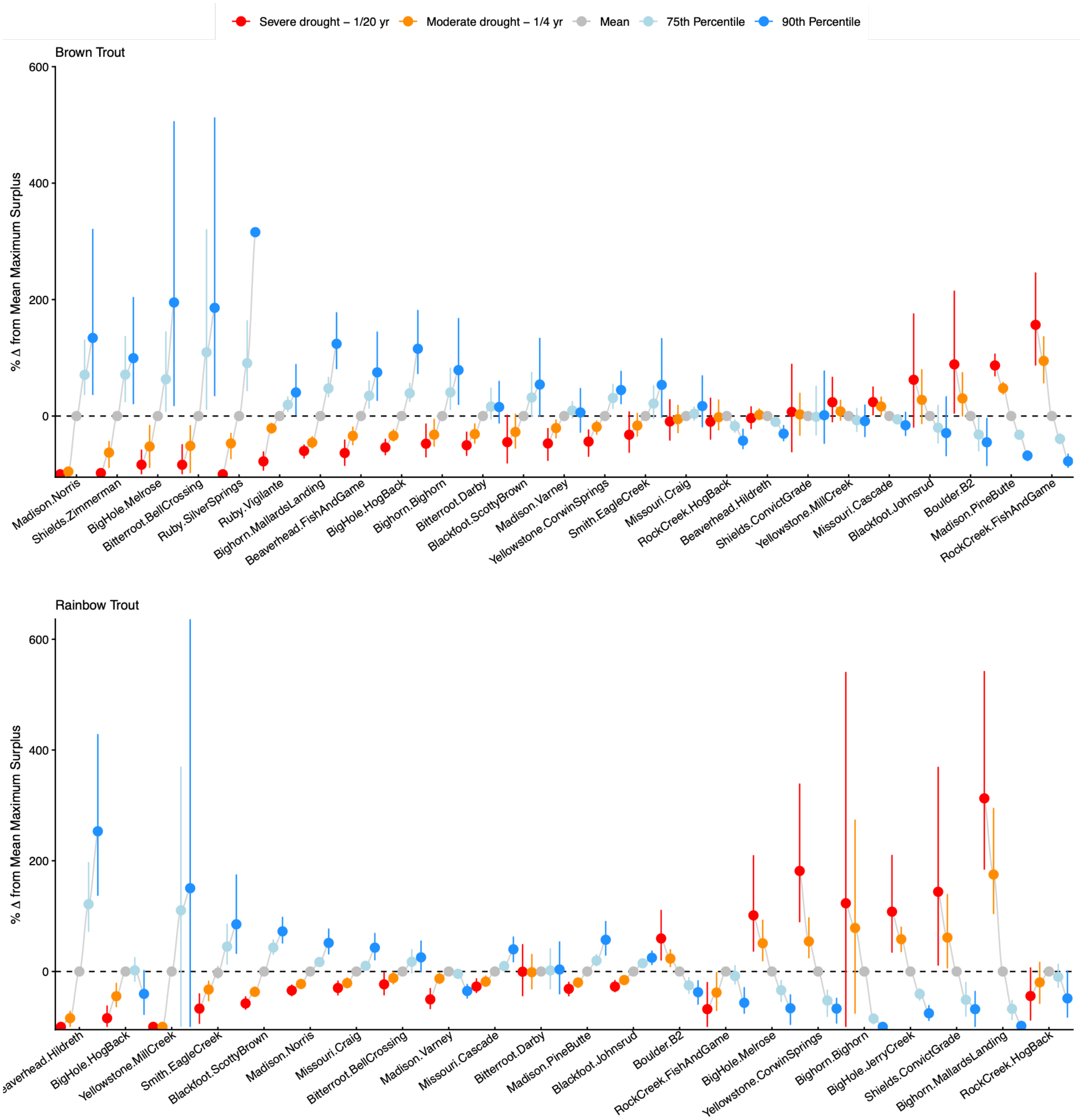


Fig. S2. Population-specific responses in surplus production (mean change from the long-term average flow scenario with 90% credible intervals) for brown trout (top) and rainbow trout (bottom) populations under the five flow scenarios, shown as percent. Population labels are expressed as “river.section,” indicating river identity followed by section designation.

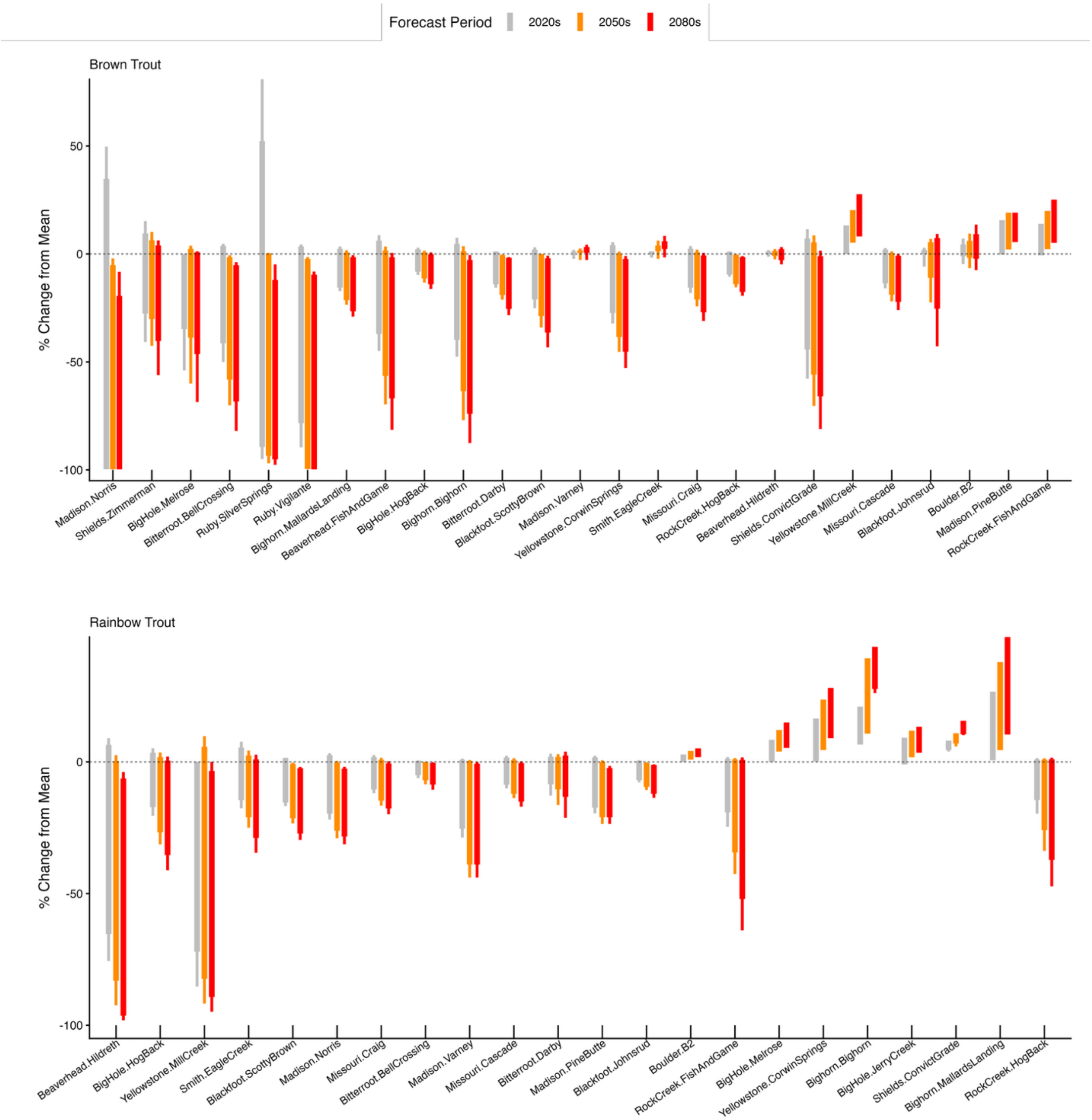


Fig. S3. Percent change in carrying capacity for each brown trout and rainbow trout population under projected changes in mean summer streamflow. Boxplots show the distribution of percent change in carrying capacity across all populations combined, with interquartile range (thick line) and 10th to 90th quantiles (thin line). Population labels are expressed as “river.section,” indicating river identity followed by section designation.
